# High-dimensional and spatial analysis reveals immune landscape dependent progression in cutaneous squamous cell carcinoma

**DOI:** 10.1101/2022.04.19.488697

**Authors:** A.L. Ferguson, A.R. Sharman, R.O. Allen, T. Ye, J.H. Lee, H. Low, S. Ch’ng, C.E. Palme, B. Ashford, M. Ranson, J.R. Clark, E. Patrick, R. Gupta, U. Palendira

## Abstract

**Purpose:** The tumour immune microenvironment impacts the biological behaviour of the tumour but its effect on clinical outcomes in head and neck cutaneous squamous cell carcinomas (HNcSCC) is largely unknown.

**Experimental Design:** We compared the immune milieu of high-risk HNcSCC that never progressed to metastasis with those that metastasised using multi-parameter imaging mass cytometry. The cohort included both immunosuppressed patients (IS) and patients with an absence of clinical immune-suppression (ACIS). Spatial analyses were used to identify cellular interactions that were associated with tumour behaviour.

**Results:** Non-progressing primary HNcSCC were characterised by higher CD8+ and CD4+ T cell responses, including numerically increased Regulatory T cells. By contrast, primary lesions from HNcSCC patients who progressed were largely devoid of T cells with lower numbers of innate immune cells and increased expression of checkpoint receptors and in the metastatic lesions were characterised by an accumulation of B cells. Spatial analysis reveals multiple cellular interactions associated with non-progressing primary tumours that were distinct in primary tumours of disease progressing patients. Cellular regional analysis of the tumour microenvironment also shows squamous cell-enriched tumour regions associated with primary non-progressing tumours.

**Conclusions:** Effective responses from both CD8+ and CD4+ T cells in the tumour microenvironment are essential for immune control of primary HNcSCC. Our findings indicate that the early events that shape the immune responses in primary tumours dictate progression and disease outcomes in HNcSCC.

**Translational Relevance:** The ability to predict metastatic tumour progression at the time of initial diagnosis of primary HNcSCC could tailor personalised medical care including disease surveillance strategies and identifying patients who will benefit most from adjuvant therapy.

**One Sentence Summary:** The immune landscape of high-risk cutaneous squamous cell carcinoma differs in tumours that never progress compared to those that progress to metastasis.

## INTRODUCTION

Squamous cell carcinoma (SCC) is the second most common skin cancer^1^. Cutaneous SCC most commonly arises in the sun exposed head and neck (HNcSCC) in Fitzpatrick skin types 1, 2 and 3 ^2,3^. Most HNcSCC can be treated with surgery with good local control. However, a subset of tumours that are large, infiltrate the subcutaneous tissue, demonstrate poor differentiation or perineural invasion (PNI) are considered to be at high-risk of local recurrence or developing metastases^1,4–6^. In clinical practice, these factors alone are unreliable as predictors of disease progression^7,8^.

Immunosuppressed patients such as organ transplant recipients are at a significantly higher risk of developing recurrent and metastatic HNcSCC ^9^ suggesting that the immune system plays a critical role in progression and control of HNcSCC. The role of the immune system in HNcSCC is further supported by data from the recent clinical trials with immune checkpoint inhibitors (ICI) that demonstrate response in 47% of patients with advanced HNcSCC^10,11^ leading to two major adjuvant immunotherapy trials in resected high-risk cSCC (NCT03969004 and NCT0383316). Thus, characterising the immune milieu and the tumour microenvironment (TME) of HNcSCC will provide insights into the immune factors that may underlie clinically aggressive behaviour and immunotherapy response in HNcSCC.

Defining the immune phenotypes within the TME has been limited and no study to date has been able to spatially resolve the immune and tumour cell interactions. Imaging mass cytometry (IMC) has recently enabled high-dimensional examination of the TME by simultaneous analyses of 36 parameters. It enables quantification of the components of the immune response and allows evaluation of spatial interactions between tumour cells, immune cells and other elements of TME^12^. Here we have applied IMC to map the immune landscape and identify differences between high-risk primary HNcSCC tumours that did not progress and those that developed metastases (progressing tumours). While most cancer registries, across the world do not collect data on HNcSCC, it is recognized that large primary high risk HNcSCC, that requires radical surgical excision, but has not progressed at long term follow up is rare^13–15^. Thus, this study not only presents data from novel technology, but also utilises a rare cohort of patients with long term follow up.

## METHODS

### Patients

Following institutional human research ethics approval, high-risk HNcSCC patients were identified from the Sydney Head and Neck Cancer Institute (SHNCI) database. High-risk was defined by the 7^th^ edition of the American Joint Commission on Cancer (AJCC) manual^16^ based on size >20mm, involvement of subcutaneous tissue, poor differentiation, or presence of perineural invasion (PNI). The formalin fixed paraffin embedded (FFPE) tissues were retrieved from the archives of the Department of Tissue Pathology and Diagnostic Oncology at the Royal Prince Alfred Hospital. All relevant clinicopathologic and follow up data were obtained from the prospectively collected SHNCI database. Clinicopathological data included age, sex, primary HNcSCC site, and immune suppression including organ transplant, non-Hodgkin’s lymphoma such as chronic lymphocytic lymphoma/leukemia (CLL) or immune suppressive drugs. Patients were classified as non-progressors (NP) if their resection demonstrated high-risk HNcSCC and the concurrent neck dissection was histologically negative for metastases. Furthermore, it was a requirement that they did not develop recurrence or metastases at a minimum follow up of 2 years. Patients were classified as disease progressors (DP) if they demonstrated nodal metastases in the concurrent neck dissection or developed metastases (regional or distant) during follow-up (Fig 1). The DP group included five immunosuppressed patients. Amongst the DP group, tissues from primary HNcSCC were available in nine patients; of these both concurrent primary and metastatic tissues were available in five patients. Tissues from only the metastases were available in the remaining 13 patients (Fig 1).

**Figure 1.**
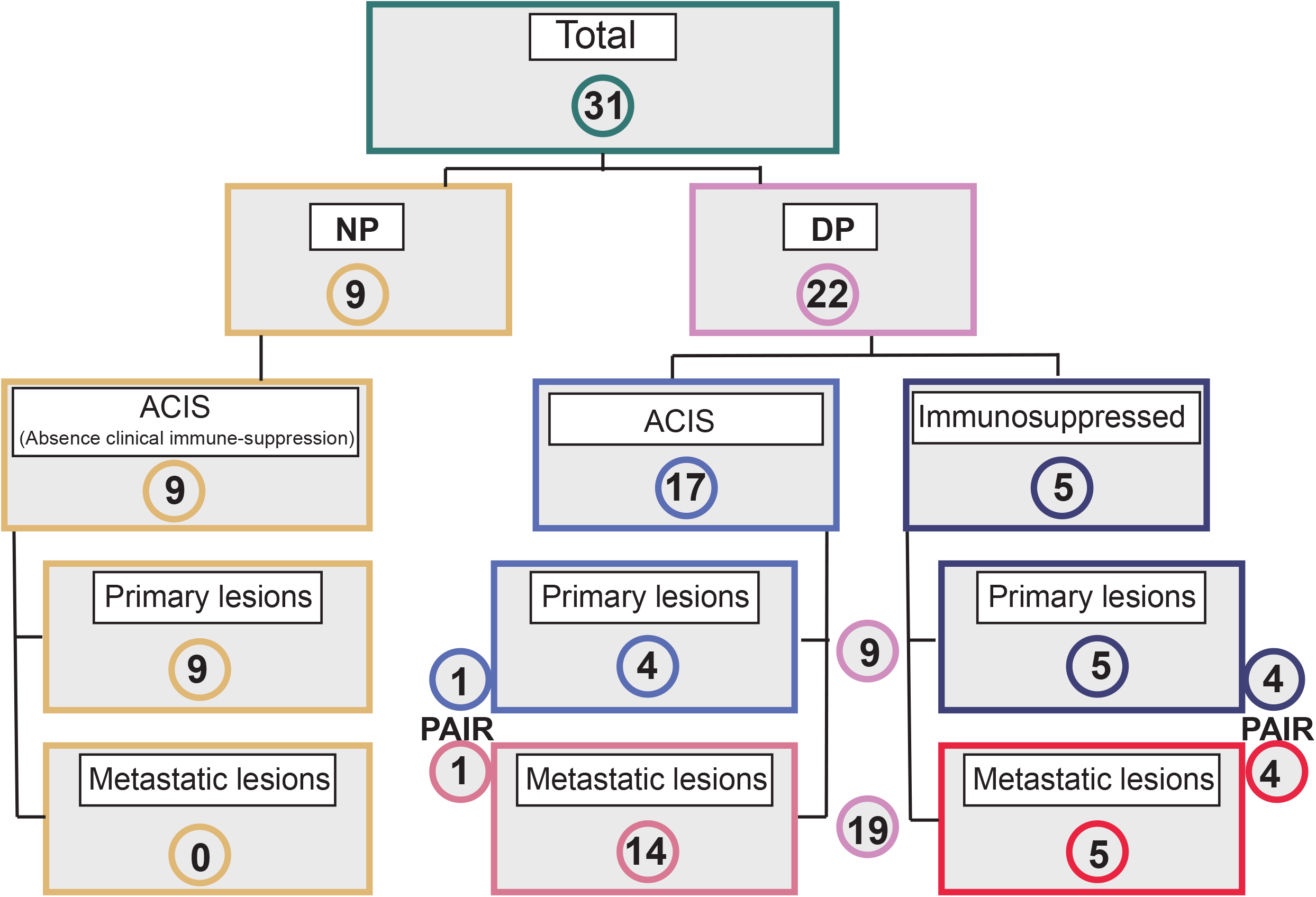
Flowchart of patient cohort, disease groups and tumour tissue classification. All patients demonstrated high-risk HNcSCC. Patients were classified as non-progressors (NP) if the concurrent neck dissection was histologically negative for metastases and they did not develop recurrence or metastases at a follow up of 2 years. Patients were classified as disease progressors (DP) if they demonstrated nodal metastases in the concurrent neck dissection or developed recurrence (local, regional, or distant) during follow-up. The DP group included five immunosuppressed (IS) patients. Amongst the DP group, tissues from primary HNcSCC tumours were available in nine patients, 5 IS and 4 patients with an absence of clinical immune-suppression (ACIS); Paired (concurrent) primary and metastatic tissues were available in five patients, 1 IC, 4 IS. Tissues from only locoregional metastases were available in the remaining 13 DP patients.

### Antibody Panel

An antibody panel was designed to identify squamous cells, cancer-associated fibroblasts, immune cells, immune checkpoint inhibitors, cell-signalling pathways and TME structural components. Clone information is available in Table 2.

### Preparation and staining

The haematoxylin and eosin stained sections of all primaries and metastases available were reviewed by head and neck pathologists and three tumoural regions of interest were selected for each case to account for heterogeneity in the TME if any. The regions included intratumoural area, the infiltrative front of the tumour and foci of perineural or lymphovascular involvement if present. Three tissue microarrays (TMAs) were created from selected regions using 1mm^2^ sample cores (Fig 2A). All three TMAs were sectioned at 7μm onto charged slides. TMA slides were deparaffinised with xylene (10 min) and rehydrated through a graded series of ethanol solutions (100, 90%; 70% and 50%) then washed in TBS-T (1x TBS with 0.05% Tween_20_). Antigen retrieval was conducted in Tris-EDTA buffer at pH 9 in a microwave. Buffer was brought to boiling point (at 100% power) followed by an additional 15 min at 20% power. The TMA slides were allowed to cool down to room temperature before proceeding. TMA slides were washed in TBS-T (10 min) prior to a second wash with PBS. TMA slides were blocked with Perkin Elmer_TM_ blocking buffer (Cat#ARD1001EA) for 45 min at 37°C. The TMA slides were then incubated overnight with antibody cocktail (refer to table 2) at 4 °C in humidity chamber. TMA slides were then washed twice in 0.1% Triton-X in DPBS and twice with PBS. TMA slides were stained with Ir-Intercalator (1:400 in DPBS, Cell-ID Fluidigm) for 30 min at RT. TMA slides were then washed in deionized H_2_O for 5 min before being allowed to air-dry at RT.

**Figure 2.**
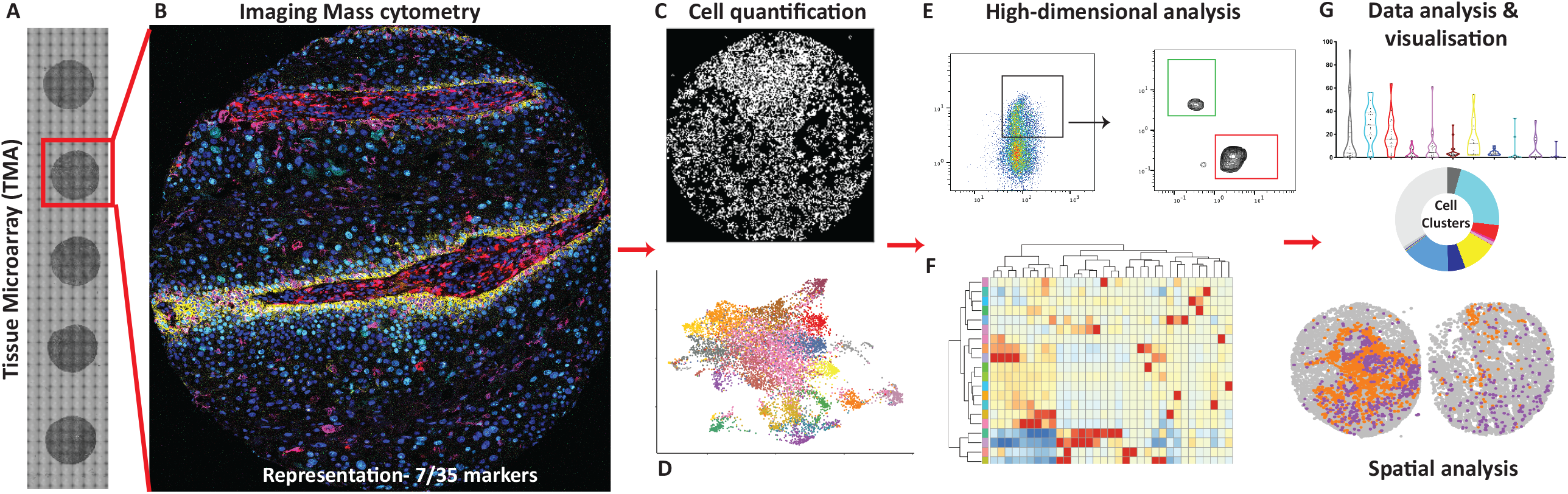
Imaging Mass Cytometry Analysis Workflow. TMAs **(A)** of human biobanked FFPE tumour tissue is stained with up to 36 metal-tagged antibodies to markers of tissue structure and cell phenotypic and functional markers, 7/36 of these markers are represented **(B**) in a human SCC tumour sample. This data is then processed using specialised software **(C)** to segment the tissue into cells (CellProfiler) and extracted as single cell data including spatial location and antibody intensities. Cells are then identified and quantified (FlowJo) **(E)**. Map using Uniform Manifold Approximation and Projection (UMAP) for dimension reduction of single cells from high-dimensional images of tumours coloured by sample **(D)** and cell-type cluster identifier **(G)**. Heat map showing the mean marker expression for each K-means cluster **(F)**. Clusters are identified, further quantified (FlowJo, GraphPad) and assessed for tissue (spatial) location **(G)**.

### Imaging mass cytometry

The TMA images were acquired using a Hyperion Imaging Mass cytometer (Fluidigm). A pulsed laser scanned and ablated the tissue. Metal ions were analysed by time-of-flight mass cytometry. Output data was produced as mass cytometry data files which were pre-processed and converted into single and multi-TIFF files using MCD viewer version 1.0.560.6 (Fluidigm) for further computational analysis. MCD viewer was used for visualising images, TIFF extraction and creating representative pseudo-colour images (Fig 2B). Cell segmentation analysis of IMC data was performed using open source CellProfiler cell image analysis software version CellProfiler 2.2.0 (Fig 2C) based on marker intensity and initial nuclear segmentation (DNA-intercolator). CellProfiler output as .csv files was converted to .fcs files using R. Quantitative analysis of IMC data was performed using FlowJo software (TreeStar) (Fig 2E). Extracted FlowJo data was analysed using GraphPad Prism version 9.0.0 (Fig 2G). Statistical analysis was performed using GraphPad 9 (Prism). Mann-Whitney statistical tests and t-tests were applied to determine significance between clinically distinct groups.

***Spatial analysis*** of the IMC data was performed in R v4.1.2. Marker intensities for each cell were calculated as the average marker intensity within the segmented nuclei. Marker intensities were normalised between each image by dividing by a trimmed mean of the marker intensity within an image, thresholding the intensities at the 99^th^ quantile and scaling to have a maximum value of one. K-means clustering with 20 clusters was used to identify cell types. A Uniform Manifold Approximation and Projection (UMAP) plot was calculated using the default settings in the UMAP package^17^ in R. The pheatmap package^18^ was used to create a heatmap of the standardised average marker intensities for each cell type, thresholded at plus and minus two. Tests for a change in spatial localisation between two cell types was performed using the spicyR package^19^ from Bioconductor. The lisaClust package^20^ (XXX) was used to classify the tissue into four distinct TME regions using a range of radii from 10 to 50 microns. Where appropriate, the p-values were adjusted for multiple comparisons, false discovery rate (FDR), using a Benjamini-Hochberg correction.

## RESULTS

### Cohort

The cohort included 31 patients with a median age of 70 years and 25 (81%) were males (Table 1, Fig 1). At a minimum follow-up of 2 years (range, 24-88 months), nine patients with primary HNcSCC never progressed (NP) and were alive without disease. Twenty-two patients presented with nodal metastases or developed metastases on follow-up (DP), including one patient who developed distant metastases and died of disease (Fig 1).

**Table 1.** Clinical Characteristics.

|  | Non-Progressors, NP<br>(n= 9) | Progression to<br>Metastases, DP<br>(n= 22) |  |
| --- | --- | --- | --- |
| Males | 6 (67%) | 19 (86%) | (overall=<br>81%) |
| Females | 3 (33%) | 3 (14%) | (overall=<br>19%) |
| Median Age (years) | 63 (39-84) | 74 (32-88) | $P=0.1120^{\wedge}$ |
| Primary Site: |  |  |  |
| Scalp | 1 | 5 |  |
| Forehead | 3 | 3 |  |
| Ear | 1 | 3 |  |
| Nose | 1 | 2 |  |
| Cheek | 2 | 2 |  |
| Lip | 1 | 2 |  |
| unknown | N/A | 5 |  |
| Median Tumour Size<br>(mm) | 25 (7-65) | Primary= 43 (23-55)*<br>Metastasis= 39.5 (14-67)* | $P=0.4847^{\#}$ |
| Immunosuppression | 0 | 5 |  |
| Treatment: |  |  |  |
| Surgery only | 4 | 3 |  |
| Surgery +<br>Radiotherapy | 5 | 19 |  |

^^^unpaired t-test with Welch’s correction. ^#^ANOVA. ^*^Data not available for all patients.

### Non-progressing primary tumours are characterised by active CD4+ and CD8+ T cell responses

To understand the immune mediated mechanisms that are critical for early control of tumours, we compared the immune landscape of NP and primary HNcSCC of DP. Focusing on the adaptive immune system, CD8+ T cells, which are critical for tumour control were greater in the NP group (p=0.006, Fig 3A I). These CD8+ T cells were also more proliferative (p<0.0001, Fig 3A II) and expressed a greater (p<0.0001) proportion of granzyme B (Fig 3A III) in the NP group.

**Figure 3.**
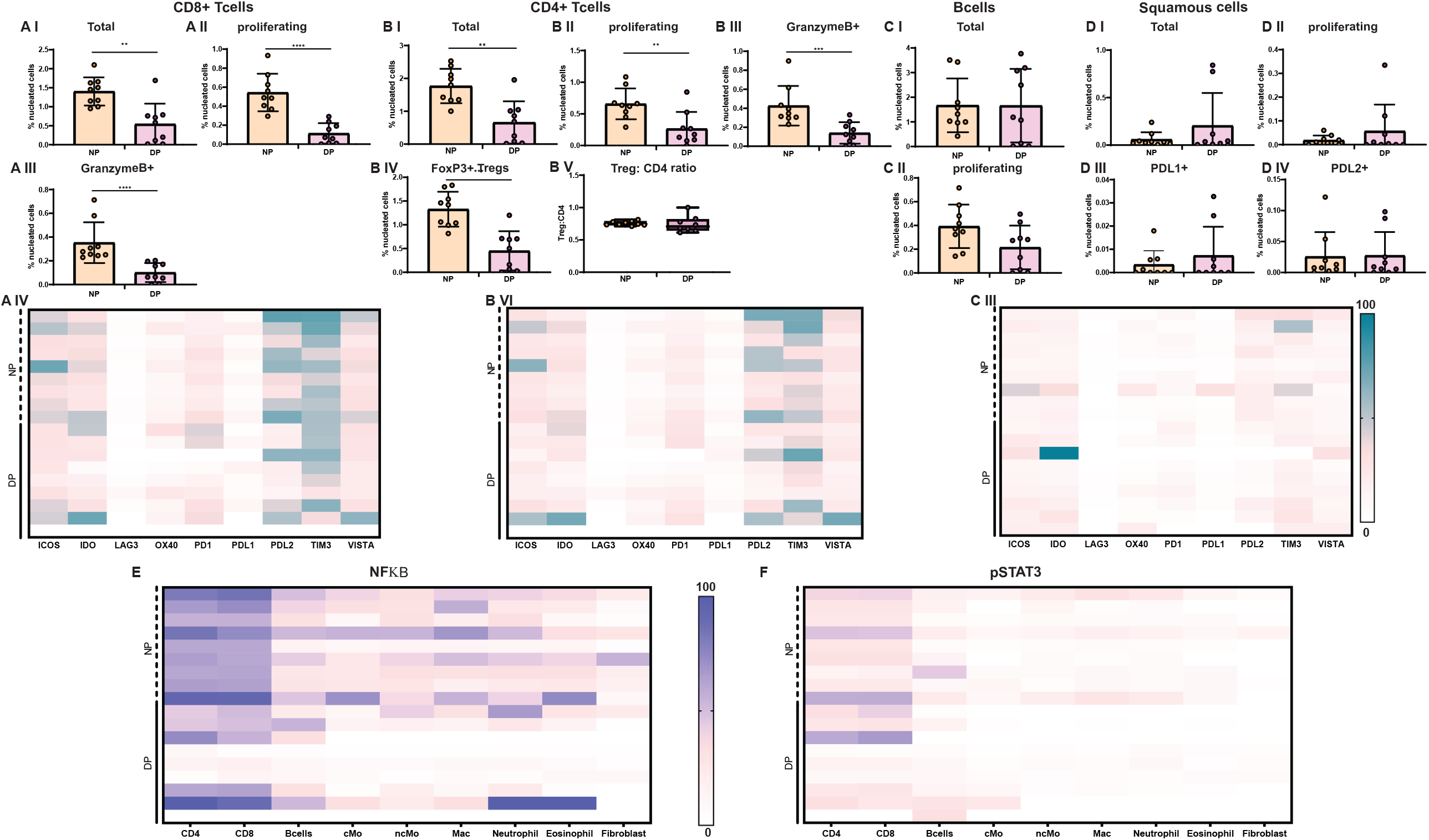
Lymphocyte infiltration and tumour cell subsets in primary cutaneous squamous cell carcinomas. **A** (**I**) Total (**II**) Proliferating and (**III**) Granzyme B expressing CD8 T cells of non-progressors (NP) n=9 and patients who progressed (DP) n=9. **(IV)** Heatmap displaying mean expression of checkpoint receptors and regulatory markers on CD8 T cells. **B** (**I**) Total **(II)** Proliferating **(III)** Granzyme B expressing CD4 T cells **(IV)** Regulatory T cells and **(V)** Treg: CD4 T cell ratio of non-progressors (NP) and Progressors (DP). **(VI)** Heatmap displaying mean expression of checkpoint receptors and regulatory markers on CD4 T cells. **C (I)** Total and **(II)** Proliferating B cells of non-progressors (NP) and progressors (DP). **(IV)** Heatmap displaying mean expression of checkpoint receptors and regulatory markers on B cells. **D (I)** Total **(II)** Proliferating **(III)** PDL1+ and **(IV)** PDL2+ squamous cells of non-progressors (NP) and progressors (DP). **E** NFkB expression and **F** pSTAT3 expression on Lymphocytes, myeloid cells, granulocytes and Fibroblasts. Mann-Whitney statistical test was applied to determine significance. *p < 0.05, ** p < 0.01, ***p < 0.001, **** p < 0.0001.

We next examined the expression of immune checkpoint receptors and co-stimulatory receptors on CD8+ T cells to determine whether dysregulated responses could also be a factor in tumour progression. The expression patterns of ICOS and OX40, two co-stimulatory molecules associated with effective T cell responses, did not show any difference between groups (Fig 3A IV). In addition, there was no difference in the expression of checkpoint receptors such as Indoleamine 2,3 Dioxygenase (IDO), OX40, LAG3, PD-1, PDL-1, TIM3 and VISTA (Fig 3A IV) amongst NP and primary HNcSCC of DP.

The number of CD4+ T cells (p=0.004) and FoxP3+ Tregs (p=0.001) were higher in the NP group (Fig 3B I and IV, respectively). Importantly, the CD4+ T cells were largely Ki67+, indicating there were more proliferating CD4+ T cells in the NP group (p=0.008, Fig 3B II). Additionally, there was no difference in the Treg: CD4 T cell ratio between NP and DP groups (Fig 3B V). The CD4+ T cells expressed granzyme B, suggesting cytotoxic potential, with higher expression found in the NP group (p=0.002, Fig 3B III). There was no difference in the expression pattern of co-stimulatory or inhibitory receptors on CD4+ T cells between the two groups (Fig 3B, VI).

There were no differences in total and proliferating B lymphocyte numbers nor the expression pattern of checkpoint receptors on B cells between the two groups (Fig 3C I, II, III). The proportion of total and proliferating squamous cells (Fig 3D I, II) and squamous cells expressing PD-L1 and PD-L2 (Fig 3D III, IV) were not significantly different (p>0.99, p= 0.71, p=0.80, and p>0.99, respectively).

Further characterisation of immune cells revealed expression of pro-inflammatory transcription factor NFkB on CD8+ and CD4+ T cells, with higher expression observed in the NP group (Fig 3E) as compared with the primary HNcSCC of DP. There was no clear pattern in the expression of phosphorylated STAT-3 levels, an indicator of response to multiple cytokines, although its expression levels were higher on T cells when compared to other immune cells (Fig 3F). Together our data suggests that active T cell responses in primary tumours are vital in preventing disease progression and their numbers may inversely predict the adverse biologic potential of the primary HNcSCC.

### There are no differences in the innate immune landscape between NP and DP Primary lesions in HNcSCC

The innate immune landscape of primary tumours of NP and DP was analysed, as early recruitment of innate immune cells could impact the immune milieu of the TME. In our cohort, the densities of innate populations were nearly 10-fold lower when compared to T and B lymphocyte populations (Figs 3 and 4). Interestingly, there was no difference in the number of monocytes (Fig 4A), non-classical monocytes (Fig 4B) and macrophages (Fig 4C) between groups. There was no difference in the expression patterns of checkpoint receptor ligands such as PD-L1 and PD-L2 or VISTA on these innate populations (Fig 4A II, B II, C II). There were no differences in the numbers of neutrophils (Fig 4D I), granulocytes (Fig 4E I) and fibroblasts (Fig 4F I) or the expression of checkpoint receptors/ligands between the primary lesions of NP and DP HNSCC (Figs 4DII, 4E II, 4F II).

**Figure 4.**
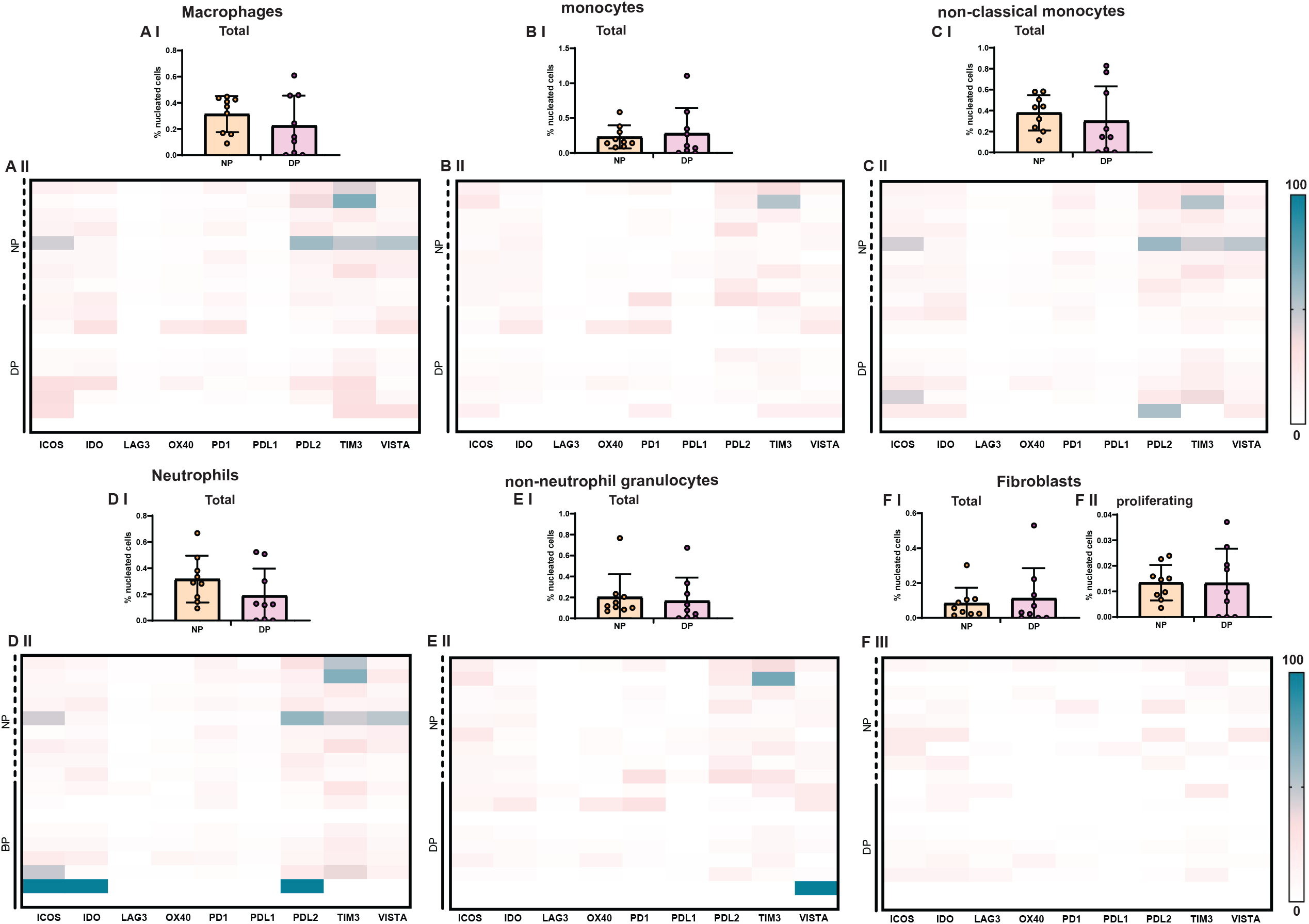
Innate immune cell infiltration and cancer-associated fibroblasts subsets in primary cutaneous squamous cell carcinomas. **A** (**I**) Total macrophages of non-progressors (NP) and progressors (DP) **(II)** Heatmap displaying mean expression of checkpoint receptors and regulatory markers on macrophages. **B** (**I**) Total monocytes of non-progressors (NP) and progressors (DP) **(II)** Heatmap displaying mean expression of checkpoint receptors and regulatory markers on monocytes. **C** (**I**) Total non-classical monocytes of non-progressors (NP) and progressors (DP) **(II)** Heatmap displaying mean expression of checkpoint receptors and regulatory markers on non-classical monocytes. **D** (**I**) Total neutrophils of non-progressors (NP) and Progressors (DP) **(II)** Heatmap displaying mean expression of checkpoint receptors and regulatory markers on Neutrophils. **E (I)** Total non-neutrophil granulocytes of non-progressors (NP) and progressors (DP) **(II)** Heatmap displaying mean expression of checkpoint receptors and regulatory markers on non-neutrophil granulocytes **F** (**I**) Total (**II**) Proliferating and Fibroblasts of non-progressors (NP) n=9 and patients who progressed (DP) n=9. **(III)** Heatmap displaying mean expression of checkpoint receptors and regulatory markers on Fibroblasts. Mann-Whitney statistical test was applied to determine significance. *p < 0.05, ** p < 0.01, ***p < 0.001, **** p < 0.0001.

### Spatial analysis reveals specific cellular interactions associated with tumour behaviour

In addition to cell densities, cellular interactions can also play a crucial role in tumour control. We therefore determined which cellular interactions within the primary tumours were associated with metastases (DP). Unbiased clustering analysis of immune populations from the IMC data sets was performed, generating 20 distinct clusters (Fig 5A) with different immune phenotypes (Fig 5B), revealing all major innate and adaptive immune populations. We then determined whether specific interactions between cell types could also differentiate NP and DP tumours. We determined which clusters interacted with each other and which clusters avoided each other (Fig 5C). Focusing on interactions or avoidance that were significantly different between NP and DP, we could identify specific interactions or avoidance that could be associated with clinical outcomes.

**Figure 5.**
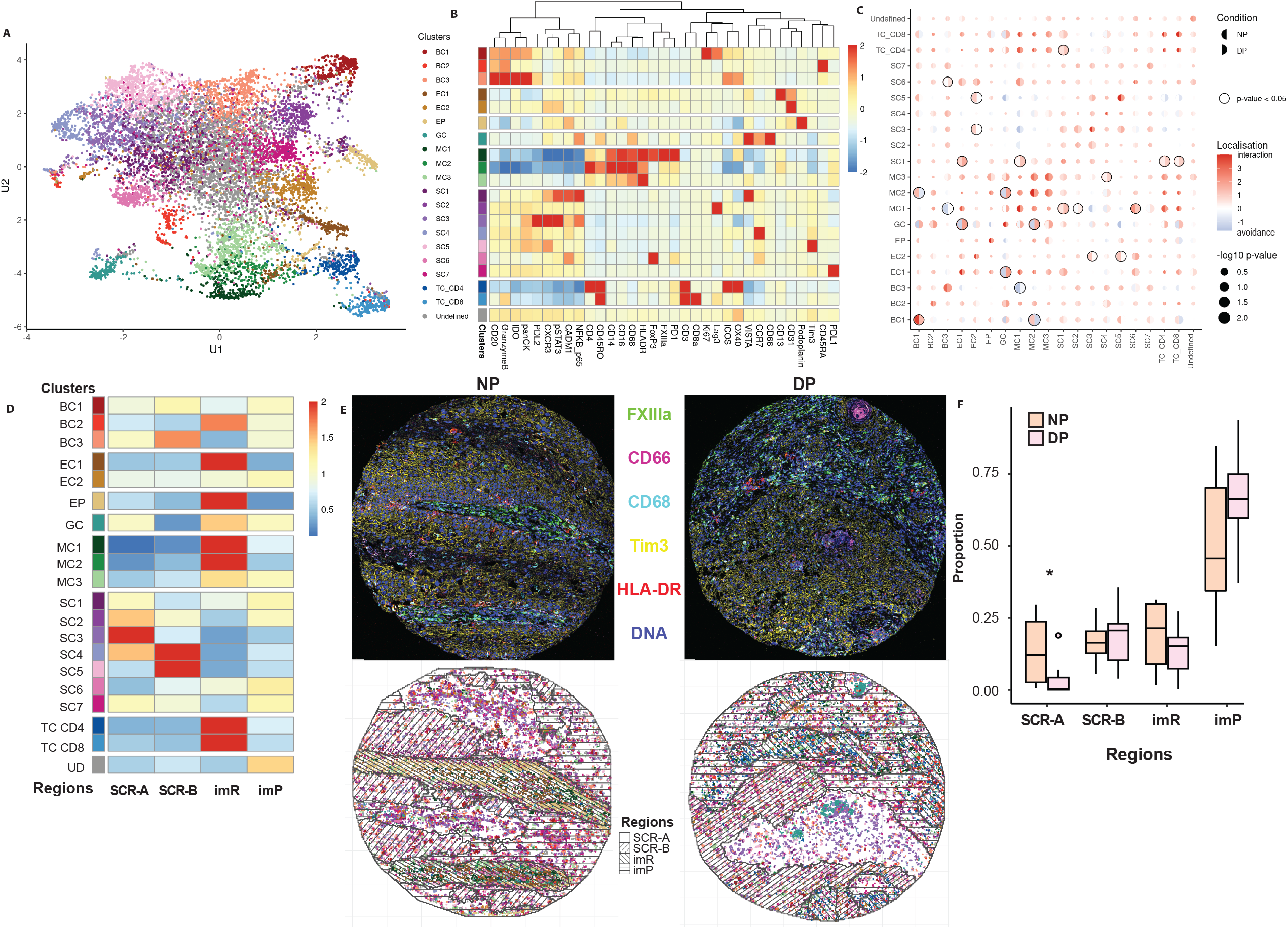
Spatial Analysis of the immune landscape of Non-progressors and Progressors in HNcSCC. **A** Representation of cell clusters identified using UMAP analysis of primary tumour samples from NP (n=9), DP (n=5) with clusters indicated by colour visualised in **B** heatmap of mean marker expression for each K-means cluster. Cells are phenotyped by their marker expression as B cell (BC), Endothelial cell (EC), Epithelial cell (SP), Granulocyte (GC), Squamous cell (SC), T cell (TC) CD4 or CD8 and undefined. **C** Spatial interactions (red) or avoidance (blue) in clusters of NP 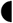 and DP 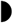 primary tumour samples as identified by spicyR package (Bioconductor), ○ p value <0.05, circle size indicates magnitude of statistical significance. **D** Four specific TME regions were identified within the tumour tissue of NP and DP patient primary tumours based on magnitude and spatial placement of cell clusters. Regions were classified as squamous cell region A (SCR-A) □, squamous cell region B (SCR-B) 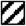, immune-milieu Rich (imR)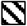 and immune-milieu Poor (imP)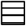 based on characteristics of cells present, represented as a heatmap **E** Representative images of NP and DP primary tumour tissue from IMC displaying 6/35 markers, FXIIIa= green, CD66= magenta, CD68= cyan, Tim3= yellow, HLA-DR= red, DNA intercalator= blue; with matching regional maps displaying cell clusters (by colour) and regions Type I SCC, Type II SCC, Cell-infiltrated and Cell-Barren. **F** Quantification of regions present in NP and DP, statistically significant difference *p<0.05.

Amongst the significant interactions associated with favourable outcomes (NP) were a subset of B cells (BC1) - Myeloid cells (MC2) interactions, presence of endothelial cells amongst tumour cells and a subset of myeloid cells (MC1) interacting with a subset of tumour cells (SC6). Interestingly, a distinction was found in that one of these interactions, BC1-MC2 was found to avoid each other in patients with poor outcomes (DP). By contrast, in patients with poor outcomes (DP), cell phenotype with features of a granulocyte (GC) interacted strongly with endothelial cells and MC2 populations, with clear avoidance between these clusters in the NP group, marking it as a potential polymorphonuclear-myeloid-derived suppressor cell

(PMN-MDSC). These interactions may point to yet unidentified roles for these immune cells in enhancing or regulating the immune milieu of the TME.

We then focussed on determining whether immune cells preferentially localised within the TME or were randomly distributed. To examine this, we identified regions based on cell clustering within the TME resulting in 4 proportion-based regions within all TMEs examined. Two of the regions were rich on tumour cell types, namely squamous cell region (SCR)-A characterised by squamous cells of immune response enhancing profile and SCR-B, differentiated by squamous cells with immune response dysregulating profile. The other two regions were either immune milieu rich (imR) or immune milieu poor (imP) (Fig 5D & E). When we compared the relative abundance of these regions across NP and DP patients, the only difference was within the distribution of SCR-A, which was higher in NP when compared to DP. These data suggest that immune cells cluster within TME in specific topography and that cell and regional interactions could both contribute to outcomes.

### Immune competency of patients defines the immune landscape of primary tumours

We separated our cohort into immunosuppressed and ACIS patients and examined the difference in adaptive immune populations. All immunosuppressed patients were in the DP group (n=5 primaries (P), n=5 metastases (M)). ACIS patients in the DP group (n=4 P, 14 M) had tumours characterised by extremely low numbers of T and B cells (Fig 6A, B & C) which were largely non-proliferative (Fig 6 AII, BII and CII) and the T cells lacking the expression of effector molecules such as granzyme B (Fig 6A III & 6B III). In addition, there was no difference in total and proliferating tumour cells and PD-L1 and PD-L2 expressing tumour cells in the NP, ACIS DP, and immunosuppressed DP patients (p=0.73, 0.34, 0.64, and 0.67, respectively. Fig 6 D I, DII, DIII, DIV).

**Figure 6.**
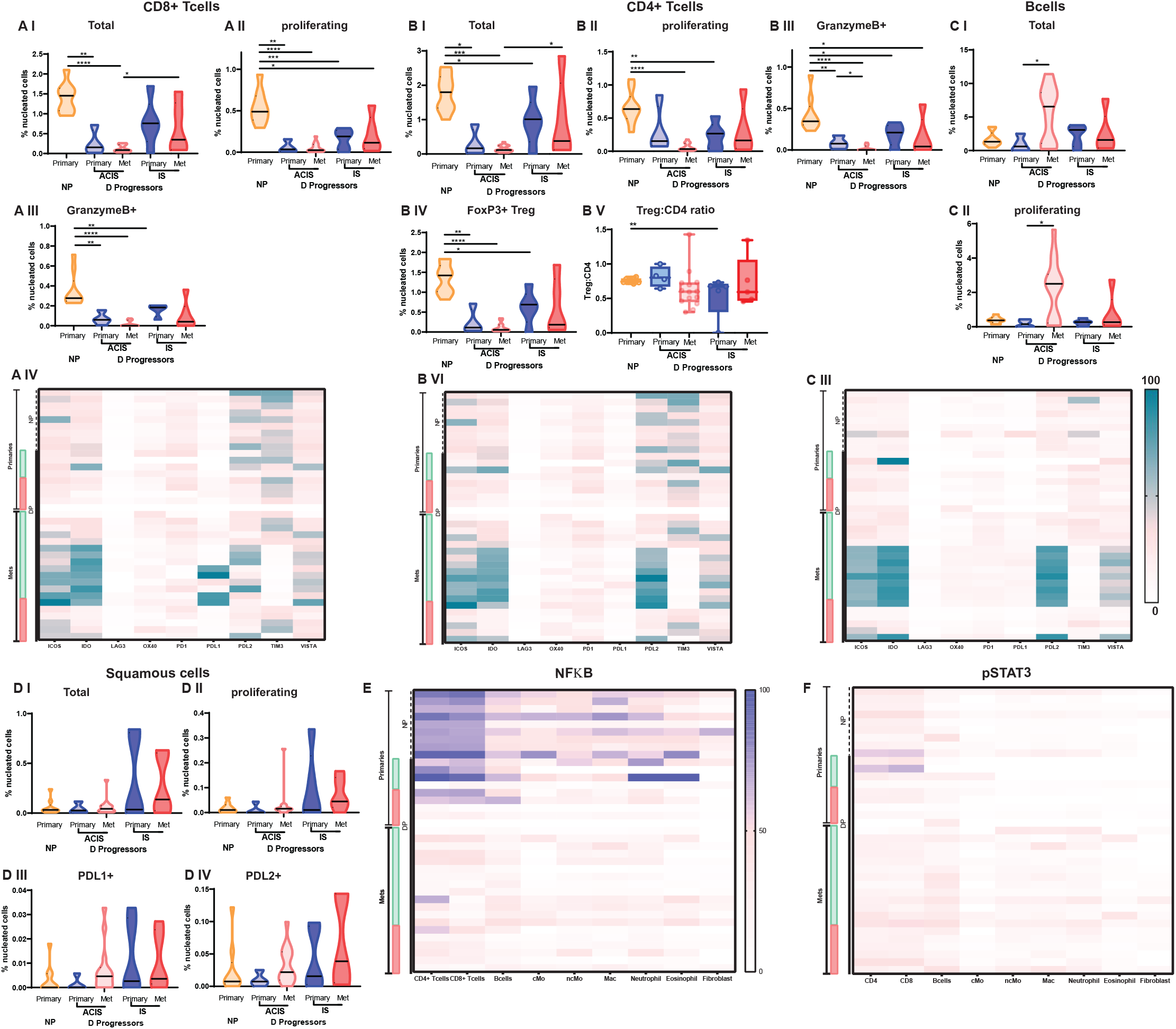
Lymphocyte infiltration and tumour cell subsets in primary and metastatic tumour samples is dependent on progression and host immune-competency. **A** (**I**) Total (**II**) Proliferating and (**III**) Granzyme B expressing CD8 T cells from primary tumours of patients that did not progress (NP) (orange), Primary tumours (pale blue) and metastases (pale red) of patients absent of clinical immune-suppression (ACIS) who progressed (DP), and primary tumours (dark blue) and metastases (red) of immunosuppressed (IS) patients who progressed (P). **(IV)** Heatmap displaying mean expression of checkpoint receptors and regulatory markers on CD8 T cells. Progressors (DP) that have an absence of clinical immune-suppression (ACIS) are represented by green bars, Progressors (P) that are immunosuppressed are represented by red bars. **B** (**I**) Total **(II)** Proliferating **(III)** Granzyme B expressing CD4 T cells and **(IV)** FoxP3+ Regulatory T cells from primary tumours of patients that did not progress (NP) (orange), Primary tumours (pale blue) and metastases (pale red) of patients absent of clinical immune-suppression (ACIS) who progressed (DP), and primary tumours (dark blue) and metastases (red) of immunosuppressed (IS) patients who progressed (P). **(IV)** Heatmap displaying mean expression of checkpoint receptors and regulatory markers on CD4 T cells. Progressors (DP) that are absent of clinical immune-suppression (ACIS) are represented by green bars, Progressors (DP) that are immunosuppressed are represented by red bars. **C (I)** Total and **(II)** Proliferating B cells of cells from primary tumours of patients that did not progress (NP) (orange), Primary tumours (pale blue) and metastases (pale red) of patients with an absence of clinical immune-suppression (ACIS) patients who progressed (DP), and primary tumours (dark blue) and metastases (red) of immunosuppressed (IS) patients who progressed (DP). **(III)** Heatmap displaying mean expression of checkpoint receptors and regulatory markers on B cells. Progressors (DP) that are absent of clinical immune-suppression (ACIS) are represented by green bars, Progressors (DP) that are immunosuppressed are represented by red bars. **D (I)** Total **(II)** Proliferating **(III)** PDL1+ and **(IV)** PDL2+ squamous cells from primary tumours of patients that did not progress (NP) (orange), Primary tumours (pale blue) and metastases (pale red) of patients absent of clinical immune-suppression (ACIS) who progressed (DP), and primary tumours (dark blue) and metastases (red) of immunosuppressed (IS) patients who progressed (DP). Heatmaps showing **E** NFКBand **F** pSTAT3 mean expression on Lymphocytes, myeloid cells, granulocytes and fibroblasts. Progressors (DP) that have an absence of clinical immune-suppression (ACIS) are represented by green bars, Progressors (DP) that are immunosuppressed are represented by red bars. Mann-Whitney statistical test was applied to determine significance. *p < 0.05, ** p < 0.01, ***p < 0.001, **** p < 0.0001.

CD4 and CD8 T cell numbers in the primary tumours of the ACIS DP group were significantly lower than the NP group (p=0.003 and p=0.003, respectively. Fig 6A & B). There were no differences in the expression patterns of co-stimulatory or inhibitory receptors expressed on T and B cells between ACIS and immunosuppressed DP patients (Fig 6 A IV, 6B V and 6C III). The primary tumours of ACIS NP patients had higher numbers of innate immune cells (Fig 7), specifically macrophages (p=0.034, Fig 7A), non-classical monocytes (p=0.020, Fig 7C) and neutrophils (p=0.011, Fig 7D). There were no differences in innate immune cells in tissues from the primary or metastases of immunosuppressed patients compared to ACIS patients (Fig 7A, B, C, D, E).

**Figure 7.**
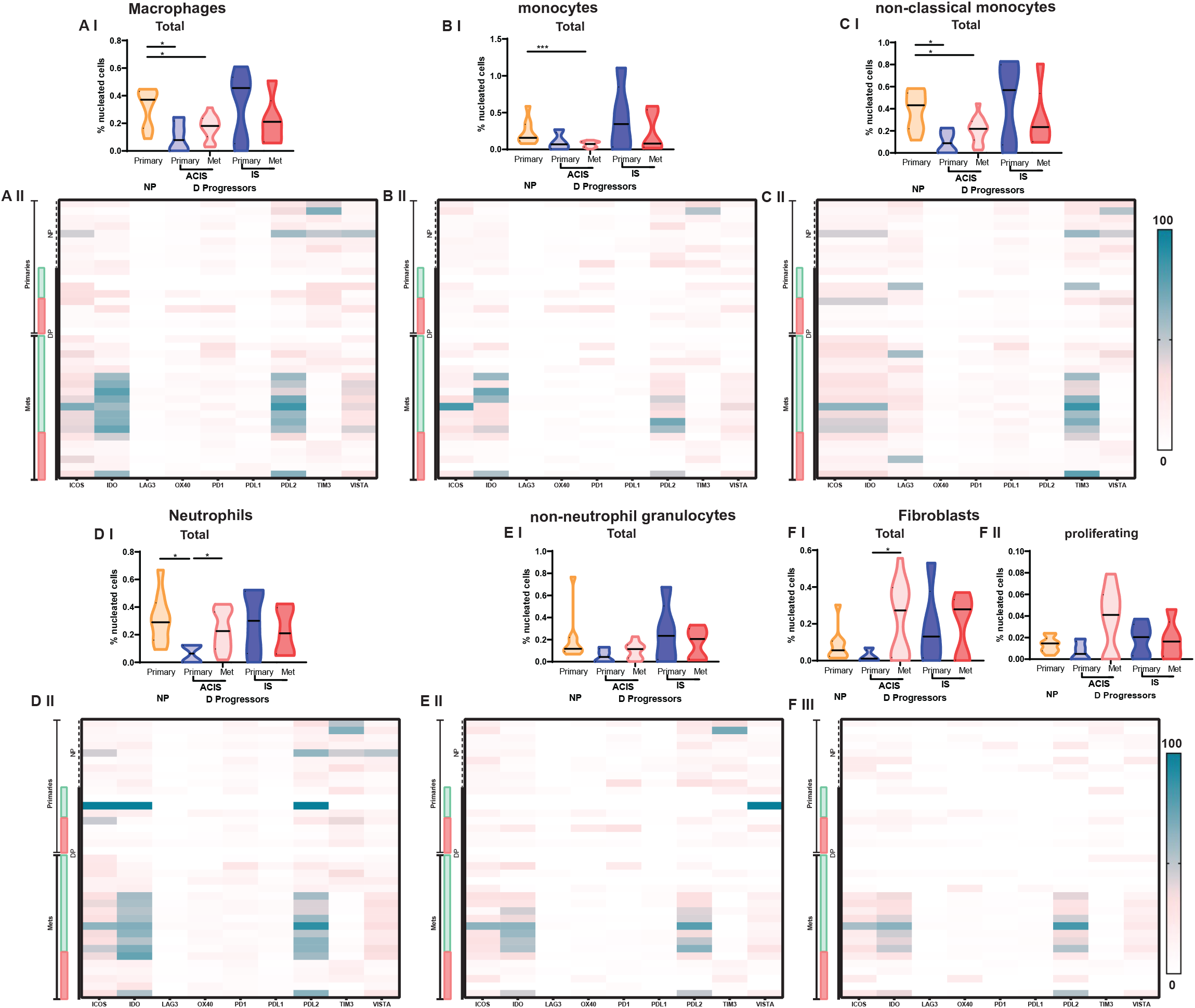
Innate immune cell infiltration and cancer associated fibroblasts in primary and metastatic tumour samples is dependent on progression and host immune-competency. **A** (**I**) Total macrophages from primary tumours of patients that did not progress (NP) (orange), Primary tumours (pale blue) and metastases (pale red) of patients with an absence of immune-suppression (ACIS) who progressed (DP), and primary tumours (dark blue) and metastases (red) of immunosuppressed (IS) patients who progressed (DP). **(II)** Heatmap displaying mean expression of checkpoint receptors and regulatory markers on Macrophages. Progressors (DP) that are absent of immune-suppression (ACIS) are represented by green bars, Progressors (DP) that are immunosuppressed are represented by red bars. **B** (**I**) Total monocytes of non-progressors (NP) and progressors (DP) **(II)** Heatmap displaying mean expression of checkpoint receptors and regulatory markers on monocytes. **C** (**I**) Total non-classical monocytes of non-progressors (NP) and progressors (DP) **(II)** Heatmap displaying mean expression of checkpoint receptors and regulatory markers on non-classical monocytes. **D** (**I**) Total neutrophils of non-progressors (NP) and Progressors (DP) **(II)** Heatmap displaying mean expression of checkpoint receptors and regulatory markers on Neutrophils. **E (I)** Total non-neutrophil granulocytes of non-progressors (NP) and progressors (DP) **(II)** Heatmap displaying mean expression of checkpoint receptors and regulatory markers on non-neutrophil granulocytes **F** (**I**) Total and **(I)** proliferating fibroblasts of non-progressors (NP) and progressors (DP) **(III)** Heatmap displaying mean expression of checkpoint receptors and regulatory markers on fibroblasts. Mann-Whitney statistical test was applied to determine significance. *p < 0.05, ** p < 0.01, ***p < 0.001, **** p < 0.0001.

### Metastatic lesions from patients with the absence of clinical immune-suppression demonstrate unfavourable immune milieu

The immune landscape in primary and metastatic tissues was compared in ACIS and immunosuppressed patients. While the T cell numbers remained consistent in ACIS patients (Fig 6AI & BI), the expression of immune checkpoints was higher in metastases, including increased expression of PD-L1 on CD8+ T cells, increased PD-L2 on CD4+ T cells and increased IDO on all T cells (Fig 6AIV & BV). This increased expression was not present in immunosuppressed patients. Interestingly, increased levels of the activating co-stimulatory checkpoint, ICOS was also observed on both CD8 and CD4+ T cells in metastatic tumours (Fig 6AIV & BV). In line with the increased expression of inhibitory receptors, the expression of NFkB and pSTAT3 were lower in T cells in metastases when compared to DP primary tumours (Fig 6D & E). By contrast, in ACIS patients there was significant increase in total B cell numbers and proliferating B cells in metastases compared to primary DP tumours (Fig 6C). This increase was not seen in immunosuppressed patients. This suggests an accumulation of dysfunctional T cells and a skewing of the immune response towards B cells within metastases of both immunosuppressed individuals and patients with ACIS.

We next explored differences in the innate immune repertoire between primary tumours and metastases. This revealed increases in neutrophils and fibroblasts in metastatic tumours from ACIS patients when compared to primary lesions (Fig 7D & F). In ACIS patients, the expression of inhibitory receptors was increased on innate populations, with the expression of IDO increased on macrophages, monocytes and neutrophils within metastatic lesions when compared to primary tumours (Fig 7 AII, BII and DII). There was also an increase in PD-L2 on macrophages and neutrophils. TIM3 was increased on non-classical monocytes in metastatic tumours of ACIS patients compared to immunosuppressed patients. These results suggest innate immune landscape of metastases is not conducive for effective immune control.

## DISCUSSION

Defining the immune mechanisms responsible for early control of tumours is critical for guiding treatment strategies. Here we show that primary HNcSCC that metastasise demonstrate a distinct immune landscape from primary tumours that do not progress, highlighting the role of the immune system in early tumour control. Using carefully curated cohorts of patients with HNcSCC with different clinical outcomes, we identified immune populations that play a critical role in the control of primary tumours and the prevention of metastasis, providing important insight into how immune control mechanisms work. Both CD4+ and CD8+ T cell numbers were significantly higher, actively proliferating, and expressed effector molecules amongst NP as compared with the primary HNcSCC tissue of DP. This difference was more apparent in ACIS patients, revealing a potential immune signature that could predict disease progression in HNcSCC. Interestingly, the malignant squamous cells with the primary tumours of NP and DP patients also showed significant differences in immune regulatory molecules expressed on their surface. The NP tumours showed significantly higher proportions of SCR-A regions. On the other hand DP tumours were more likely to show higher proportions of SCR-B regions. Our data reveals that once the tumour metastasises, the metastatic tissues harbour an unfavourable immune environment with increased expression of immune checkpoints on both T cells and innate populations. Interestingly, we also show active expansion of B cells in metastatic tissue indicating a possible regulatory role of these B cells^21–23^ which may act systemically in draining lymph nodes to promote metastasis. These findings highlight the potential for B cell-targeted immunotherapy for HNcSCC patients at high-risk of progression. B cell-targeted immunotherapy has shown potential in animal models of keratinocyte cancers^24^ to prevent development of metastases and improve survival and is currently in clinical use for treatment of leukemia and auto-immune disease. This study also highlights many potential therapeutic targets present on squamous cell populations. Antibodies targeting Lag-3 combined with anti-PD-1 are currently showing promise in clinical trials of advanced melanoma, with a progression-free survival rate at 12-months of 47.7%^25^, this regime has now begun clinical trials in other types of cancer. The presence and over-expression of Tim-3 has been shown to correlate with poor prognosis in acute myeloid leukemia (AML)^26^ and Breast cancer^27^. Whilst immune-therapeutics targeting Tim-3 have not been successful, Tim-3 is an important prognostic marker in many cancers.

This is the first study mapping both innate and adaptive immune populations of primary and metastases of HNcSCC tissues, comparing the immune landscape in patients with different clinical outcomes. One of the strengths of this study is the carefully curated cohort of HNcSCC with different clinical outcomes. While HNcSCC is one of the most common malignancies, most patients develop small lesions that are treated in the community with topical agents or excisions that may not receive comprehensive histologic examination. Large high risk HNcSCC that requires radical resection and comprehensive histologic examination but do not develop metastases at follow up, as included in this cohort, are extremely rare. Utilising this specific cohort has provided valuable insights into the characteristics of the malignant squamous cells as well as the TME of high risk HNcSCC that do not progress. This information can be utilised to predict primary HNcSCC that will likely progress by utilising the differences in the immune landscape, in particular lower numbers of T cells is useful in routine clinical practice to determine which patients will require closer surveillance and benefit most from adjuvant therapy. This is particularly evident in ACIS patients, whose tumours with low immune cell infiltration were associated with progression to metastases.

The role of cytotoxic T cells in controlling tumours is now well-established with increased CD8+ T cells being associated with good prognosis in many cancers^28–37^. In primary NP HNcSCCs, there was an increase in number, function, and proliferation of CD8+ T cells. In addition, the low expression of checkpoint receptors in the CD8+ T cell found in NP primaries, including PD-1, confirmed they were still active within the tumour microenvironment. However, the role of CD4+ T cells in tumour control is not fully understood. In our study, like their CD8+ T cell counterparts, the CD4+ T cells were likely cytotoxic and actively proliferating, suggesting a critical role for these cells in controlling primary tumours. Interestingly, the number of Tregs corresponded with increased T cell activity in NP primaries.

Importantly, our study highlights the crucial role of cellular interactions in disease progression of HNcSCC. The interaction of a specific myeloid cell (MC2) was associated with different outcomes depending on the cell it was seen interacting with. While MC2 was found to be significantly interacting with B cell subset BC1 in NP patients, these two cells showed spatial avoidance in primary tumour of DP patients. On the other hand, MC2 was seen to interact with the specific granulocyte (GC) with features of PMN-MDSCs, within HNcSCC of DP patients. A spatial avoidance between MC2 and GC was seen in NP. It is well known that B cells have the capability of regulating the development and phenotypic maturation of myeloid cells through the secretion of cytokines^38^. In the TME, this polarisation can be the difference between a myeloid cell becoming capable of recruiting T cells or evolving to a Myeloid-derived suppressor cell (MDSC) or one capable of invoking anti-tumour immune responses. One subgroup of MDSCs, PMN-MDSCs is distinct in its phenotypic identification as a granulocyte by its cell-surface expression of CD66b and lack of CD14 in humans^39,40^. Using density-gradient separation and microarray analysis in patients with lung cancer and head and neck cancer, gene-expression profiles of neutrophils and PMN-MDSCs from the same patient were found to be very different^39^. The same group found that in PMN-MDSCs, *IL-6, IFNγ* and *NF-*К*B* genes were enriched ^40^. In our study, the GC cluster identified closely resembles the signature for a PMN-MDSC cell in that it is CD14-CD66b+ and has expression of the NFКBp65 subunit. Our spatial analysis shows that this cell and it interactions with MC2 and EC2 is associated with DP patients whereas avoidance between GC and MC2 and endothelial cells is associated with NP patients. This suggests that the cell termed GC may in fact have suppressive or negative effects in the TME associated with a PMN-MDSC. This data clearly demonstrates the heterogeneity in immune cell-tumour cell interactions amongst HNcSCC with different disease outcomes. Further studies are needed to determine the biological significance of these interactions in HNcSCC.

Regional analysis is able to add extra insight into the cells present within topographical regions of the TME and their organisation within the TME of HNcSCC patients. In our study we were able to identify specific regions associated with different cellular landscapes in primary tumour from NP and DP patients. NP patients had an increased proportion of the region SCR-A which was mainly differentiated by the presence of three clusters of squamous cells SC2, SC3, SC4 and their expression of Lag 3, PD-L2, CXCR3, pSTAT3 and NFКBp65 and CCR7. This finding, particularly the role of these molecules in enhancing the immune response and control of proliferation and migration of squamous cells requires further investigation. Of note, the DP tumours lacked SCR-A regions; however showed SCR-B regions with expression of Tim-3 and FoxP3. The adverse prognostic implications of Tim-3 and FoxP3 are well established in a variety of other malignancies including AML^26^ and Breast cancer^27^. Interestingly there was no difference between NP and DP groups in the proportion of immune-milieu rich and poor regions.

The innate immune landscape was not different between primary tumours of NP and DP groups. Innate immune cells can both be beneficial or detrimental to tumour control. Tumour derived factors can impact the quantity and quality of innate cells recruited to the tumour ^41^, which in turn can impact the recruitment of other immune cells. T cell recruiting chemokines such as CXCL9 and CXCL-10 are produced by macrophages and dendritic cells, which will enable the recruitment of effector T cells that express the receptor CXCR3^42,43^. In our study, neither cell densities nor the expression pattern of immune checkpoints was different between NP and DP groups, however, it is possible that these innate cells were functionally different. It was however clear that as tumours metastasise, the innate landscape changes. Compared to their matched primaries, metastatic lesions had significantly higher proportion of neutrophils and increased expression of checkpoint receptors such as ICOS and PD-L1. This mirrors the T cell populations in metastatic lesions, suggesting that metastatic HNcSCCs have an unfavourable TME for immune control.

Immunosuppressed patients, particularly organ transplant recipients and patients with non-Hodgkins lymphoma, are up to 65 times more likely to develop HNcSCC and 5.3 times more likely to develop metastases^4^. In our cohort, the immunosuppressed patients had lower T cell responses when compared with NP group but higher than the DP group, pointing to a possible threshold effect of T cells. Despite the presence of a B cell malignancy in most of these patients, an increased B cell infiltration was not seen within the TME, highlighting alternate contributing factors facilitating poor immune control. Another possible contributing factor are innate immune cells, as the TME of immunosuppressed patients were characterised by increased numbers of monocytes, macrophages, and neutrophils, possibly contributing to poor immune control. Metastatic tumours of immunosuppressed patients showed a different immune profile from that of ACIS patients in that there was reduction in immune cell infiltrates and there were no significant changes to checkpoint receptor expression. These observations suggest that the mechanisms that underpin poor immune control of tumours are different between immunosuppressed and ACIS patients.

Mapping the immune landscape in distinct stages of HNcSCC provides valuable insights into the immune mechanisms that are effective at controlling tumour metastasis. Defining the immune composition will not only provide opportunities to develop immune signatures for predicting disease outcomes of the tumour, but also provides some basis for immune interventions. Our data suggests that primary HNcSCC that does not metastasise has a richer milieu of proliferating CD8+ and CD4+ T cells with higher expression of effector molecules as compared to tumours likely to metastasise. Further, metastatic HNcSCCs express multiple immune checkpoint receptors that could be targeted through checkpoint blockade therapies.

## Acknowledgements

This work was funded by the University of Sydney’s Sydney Research Excellence Initiative grant, Sydney Catalyst, Chris O’Brien Lifehouse, and the Cancer Institute NSW Translational Program Grant.

We acknowledge the support of the University of Sydney, Sydney Cytometry Facility.

## Author Contributions

AF, RG and UP designed the study. AF and RA performed the experiments. All authors contributed to data analysis/interpretation. AF, RG and UP wrote the first manuscript draft and all authors provided revision to the scientific content of the final manuscript.

## Figure legends

**Supplementary Figure 1**

Estimation plot showing mean differences of **A** (I) Total and (II) proliferating CD8 T cells, **B**(I) Total and (II) proliferating CD4 T cells, **C** Tregs, **D** (I) Total and (II) proliferating B cells, **E** (I) Total and (II) proliferating Fibroblasts, **F** Macrophages, **G** Monocytes, **H** Non-classical monocytes, **I** Neutrophils and **J** non-neutrophil granulocytes between matched primaries tumours and metastases from patients who progressed. Symbol ‘+’ indicates an absence of clinical immune-suppression (ACIS), ‘*’ indicates immunosuppressed patients. Estimation plot and significance generated using paired t test, significance denoted in top right corner (red *p < 0.05, ** p < 0.01).

**Supplementary Table 1. <u>IMC antibody details</u>**

