## Supplemental Table 1, Supplemental Figure 1 for "High-dimensional and spatial analysis reveals immune landscape dependent progression in cutaneous squamous cell carcinoma"

| Metal tag | Target | Antibody Clone | Company | Catalogue number |
|---|---|---|---|---|
| **139La** | pan-Cytokeratin | SPM115, SPM116 | Novus biologicals | NOVNBP234 |
| **141Pr** | CD20 | H1 | BD | 555677 |
| **142Nd** | Histone H3 | Polyclonal | Novus biologicals | NOVNB500171 |
| **143Nd** | CD45RA | HI100 | Biolegend | 304102 |
| **146Nd** | CD8a | D8A8Y | Cell signalling | 85336BF |
| **147Sm** | Podoplanin | Polyclonal | RnD Systems | RDSAF367 |
| **148Nd** | CD16 | EPR16784 | abcam | Ab256582 |
| **149Sm** | CADM1 | Polyclonal | Invitrogen | PA524196 |
| **150Nd** | IDO | D5J4E | CST | 86630S |
| **151Eu** | PDL1 | E13LN | CST | 13684S |
| **152Sm** | CD13 | 498001 | RnD Systems | MAB3815 |
| **153Eu** | CD68 | KP1 | Biolegend | 916104 |
| **154Sm** | VISTA | D1L2G | CST | 64953S |
| **155Gd** | CD31 | EPR3094 | abcam | Ab207090 |
| **156Gd** | CXCR3 | 49801 | abcam | ab64714 |
| **158Gd** | pSTAT3 | 4/P-STAT3 | Fluidigm | 3158030D |
| **159Tb** | CCR7 | Polyclonal | Proteintech | 55425-1-AP |
| **160Gd** | CD14 | EPR3653 | abcam | Ab214438 |
| **161Dy** | FXIIIa | Polyclonal | Affinity Biologicals | SAF13A-AP |
| **162Dy** | FoxP3 | Polyclonal | RnD Systems | RDSAF3240 |
| **163Dy** | PD1 | EPR4877(2) | abcam | Ab186928 |
| **164Dy** | CD45RO | UCHL1 | Becton Dickinson | 555491 |
| **165Ho** | OX40 | E9U70 | CST | 61637S |
| **166Er** | NFκB p65 subunit | K10x | Fluidigm | 3166026D |
| **167Er** | CD66a | YTH71.3 | abcam | Ab128720 |
| **168Er** | Ki67 | Polyclonal | R&D systems | RDSAF7617 |
| **169Tm** | Lag3 | D2G40 | CST | 15372 |
| **170Er** | CD3 | Polyclonal | abcam | AB5690 |
| **171Yb** | Granzyme B | EPR8260 | abcam | Ab226162 |
| **172Yb** | PDL2 | D7U8C | Fluidigm | 3172031D |
| **173Yb** | CD4 | EPR6855 | abcam | AB181724 |
| **174Yb** | HLA DR | EPR3692 | abcam | AB215985 |
| **175Lu** | ICOS | D1K2T | CST | 89601S |
| **176Yb** | TIM3 | D5D5R | CST | 45208S |
| **191Ir** | DNA intercalator | - | Fluidigm | 201192B |
| **193Ir** | DNA intercalator | - | Fluidigm | 201192B |

### Supplementary Figure 1

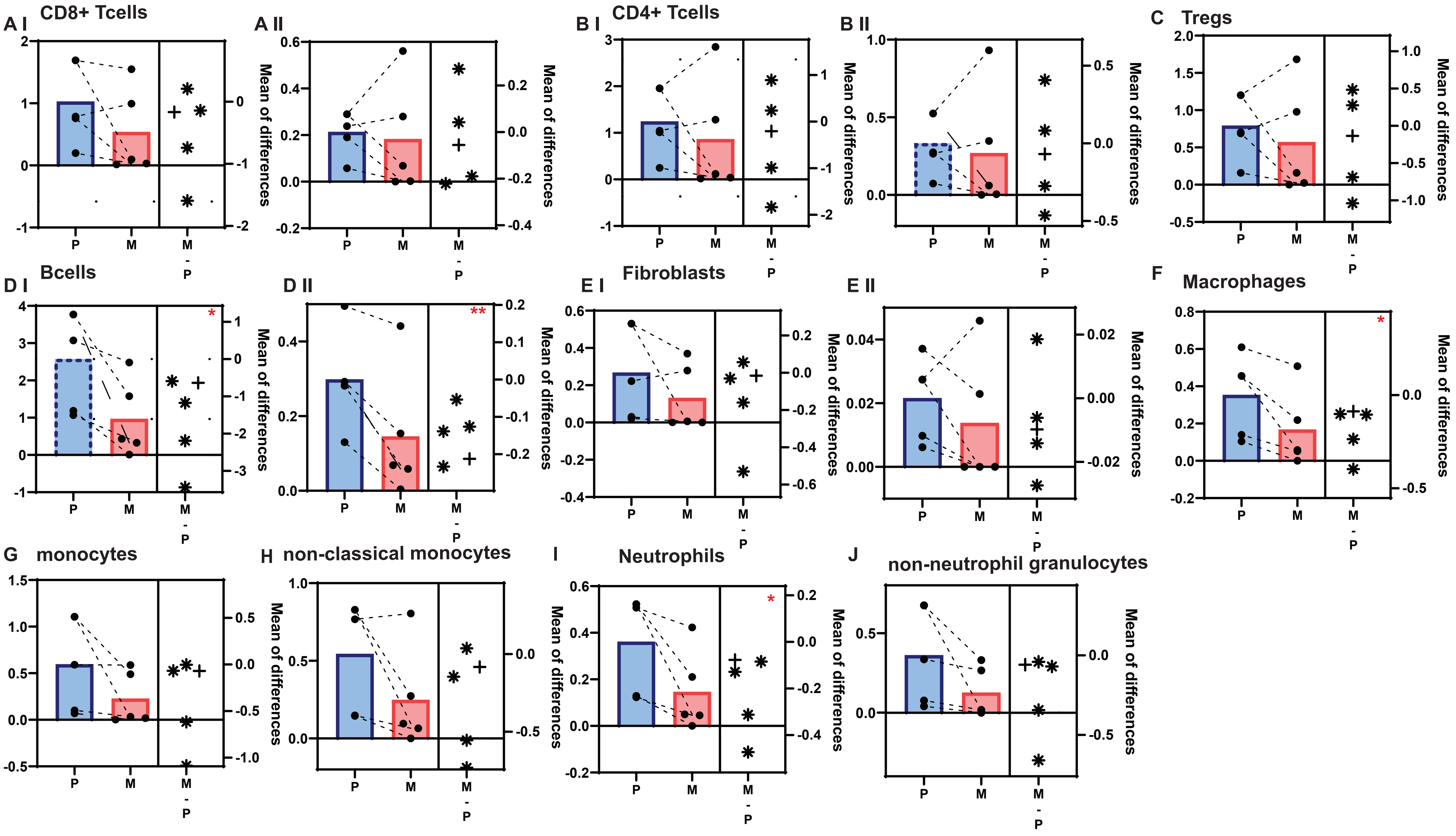
